# Prioritization of metabolic genes as novel therapeutic targets in estrogen-receptor negative breast tumors using multi-omics data and text mining

**DOI:** 10.1101/515403

**Authors:** Dinesh Kumar Barupal, Bei Gao, Jan Budczies, Brett S. Phinney, Bertrand Perroud, Carsten Denkert, Oliver Fiehn

## Abstract

Estrogen-receptor negative (ERneg) breast cancer is an aggressive breast cancer subtype in the need for new therapeutic options. We have analyzed metabolomics, proteomics and transcriptomics data for a cohort of 276 breast tumors (MetaCancer study) and nine public transcriptomics datasets using univariate statistics, meta-analysis, Reactome pathway analysis, biochemical network mapping and text mining of metabolic genes. In the MetaCancer cohort, a total of 29% metabolites, 21% proteins and 33% transcripts were significantly different (raw *p* < 0.05) between ERneg and ERpos breast tumors. In the nine public transcriptomics datasets, on average 23% of all genes were significantly different (raw *p* < 0.05). Specifically, up to 60% of the metabolic genes were significantly different (meta-analysis raw *p* < 0.05) across the transcriptomics datasets. Reactome pathway analysis of all omics showed that energy metabolism, and biosynthesis of nucleotides, amino acids, and lipids were associated with ERneg status. Text mining revealed that several significant metabolic genes and enzymes have been rarely reported to date, including PFKP, GART, PLOD1, ASS1, NUDT12, FAR1, PDE7A, FAHD1, ITPK1, SORD, HACD3, CDS2 and PDSS1. Metabolic processes associated with ERneg tumors were identified by multi-omics integration analysis of metabolomics, proteomics and transcriptomics data. Overall results suggested that TCA anaplerosis, proline biosynthesis, synthesis of complex lipids and mechanisms for recycling substrates were activated in ERneg tumors. Under-reported genes were revealed by text mining which may serve as novel candidates for drug targets in cancer therapies. The workflow presented here can also be used for other tumor types.

## Introduction

Estrogen receptor signaling is one of the main molecular features that determines the aggressiveness and the clinical course of breast cancer. Estrogen receptor negative breast tumors are aggressive and have a poor prognosis due to their high proliferation rate and their resistance to many therapeutic approaches. The estrogen-independent growth of ERneg tumors depends on a range of biological pathways, including central energy and nucleotide metabolism^33, 38^, motivating to characterize metabolic dysregulations associated with the aggressive tumor phenotype. Human metabolic network’s operation and regulation is governed by up to 10% genes in the human genome. Many of these genes and associated pathways are dysregulated and fuel a tumor’s growth, therefore they are potential drug targets.

How to identify these dysregulations in aggressive breast tumors and rank them? One of the experimental approaches is to analyze the tumors with omics assays including metabolomics, proteomics and transcriptomics. In fact, breast tumors are extensively analyzed using these assays ^1, 11, 17, 18, 22, 24, 25, 27, 29, 32, 39^. Tumor metabolomics reveals new insights into breast cancer metabolism ^6, 8, 31, 44^ MetaCancer consortium has been built to identify and validate new breast cancer biomarkers based on metabolomics^6, 7^. We have previously shown that beta-alanine and glutamate to glutamine ratio (GGR) are associated with aggressive phenotype in the MetaCancer cohort ^6, 7^. Furthermore, a multi-omics approach has been successfully used to characterize metabolic dysregulation and identify potential new therapeutic targets in lung cancer ^13^, but such investigations are limited for breast tumors. These approaches yield different lists of potential targets, creating a challenge to identify which genes and pathways can be targeted in follow-up experiments. A handful of genes are over-studied in reference to tumor biology, for example IDH1 or FH, indicating a selection bias. However, poorly-studied significant genes and associated metabolic pathways provide an additional repertoire to identify new drug targets beyond the handful of genes.

In the present study, we integrated tissue-based metabolomics with proteomics and transcriptomics in the MetaCancer cohort to identify the metabolites, proteins and metabolic genes that are associated with ERneg phenotype. To identify metabolic genes that were poorly studied in ERneg tumors, we performed Gene Expression Omnibus (GEO) based meta-analysis of nine publicly available datasets, and ranked the metabolic genes based on their significance in meta-analysis and literature count.

## Results

### Workflow to prioritize tumor metabolic genes

We applied a new bioinformatics workflow (Figure 1) to characterize metabolic dysregulations between ERpos and ERneg breast tumors. We used metabolite, protein and transcript measurements data from the MetaCancer cohort ^6, 7^ in addition to publicly available gene expression data (Table S1). The workflow incorporates significance testing, gene expression meta-analysis, pathway analysis, network mapping and text mining. The output of this workflow yields lists of metabolic pathways and under-studied metabolic genes, here associated with ERneg breast tumor biology.

**Figure 1.**
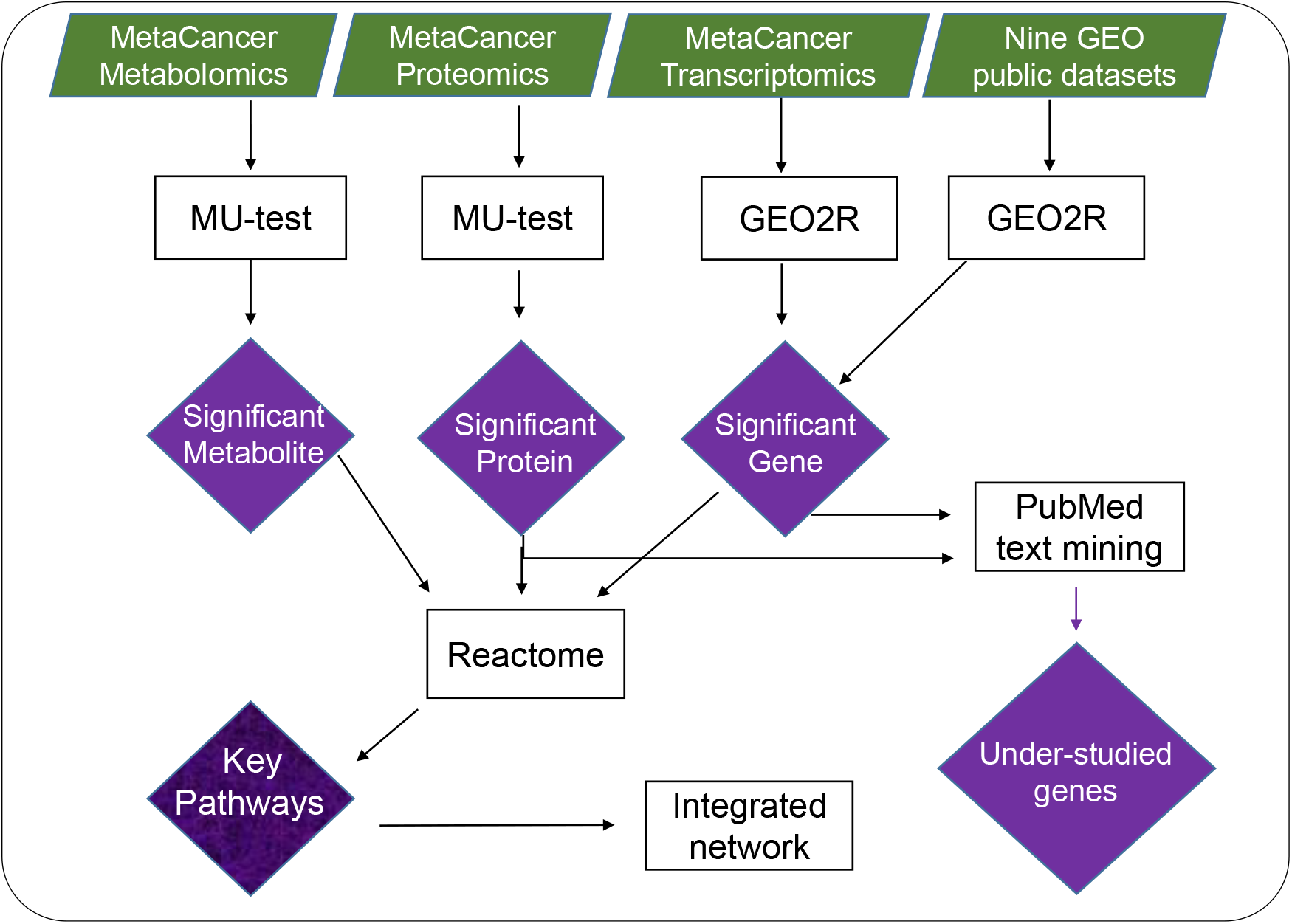
Overview of multi-omics data mining to reveal metabolic dysregulation by integrating raw p-values from metabolite, protein and gene expression analysis. Results of significance testing on individual omics-levels were subsequently analyzed by pathway enrichment analysis, by mapping to biochemical networks and by text mining. The overall outcome of such analysis is a list of key pathways and genes that includes well-studied genes as well as genes that have rarely been reported before in the context of breast cancer.

#### Molecular differences between ERneg and ERpos breast tumors

Table 1 summarizes significantly different transcripts, proteins and metabolites between ERneg and ERpos tumors in the MetaCancer and the public datasets. We used raw *p*-values to interpret the molecular differences across all multi-omics levels of cellular regulation using bioinformatics approaches. Hence, we did not correct *p*-values for multiple hypothesis testing because our objective was not to find a diagnosis biomarker panel that can distinct the tumor subtypes.

**Table 1.** Statistical results for ten GEO studies comparing ERneg vs ERpos breast tumors. Detailed results are provided in Table S6. * *indicates the MetaCancer study*

| GEO ID | entity type | ERneg Samples | ERpos Samples | Total entities | Altered entities | DOWN | UP |
| --- | --- | --- | --- | --- | --- | --- | --- |
| 59198* | transcript | 32 | 122 | 18401 | 6040 | 3103 | 2937 |
| 59198* | proteins | 29 | 96 | 1185 | 244 | 77 | 167 |
| 59198* | metabolites | 59 | 192 | 470 | 164 | 31 | 133 |
| 22093 | transcript | 56 | 42 | 22283 | 4103 | 1866 | 2237 |
| 23988 | transcript | 29 | 32 | 22283 | 5959 | 2427 | 3532 |
| 20437 | transcript | 9 | 9 | 22283 | 1191 | 653 | 538 |
| 6577 | transcript | 10 | 78 | 11171 | 4646 | 2275 | 2371 |
| 88770 | transcript | 11 | 106 | 54675 | 8043 | 3409 | 4634 |
| 22597 | transcript | 45 | 37 | 22283 | 4984 | 2528 | 2456 |
| 26639 | transcript | 88 | 138 | 54675 | 19659 | 7404 | 12255 |
| 74667 | transcript | 26 | 69 | 41093 | 7456 | 3442 | 4014 |
| 75678 | transcript | 25 | 29 | 45220 | 5958 | 3100 | 2858 |

##### MetaCancer dataset

The metabolomics data of the MetaCancer study consisted of 470 metabolites detected by gas chromatography/time of flight mass spectrometry, including 161 identified metabolites and 309 unidentified metabolites. Breast tumors from 251 women were studied, of which 192 were ERpos and 59 were ERneg. A total of 164 metabolites (63 identified and 101 unidentified metabolites) were significantly different between ERpos and ERneg cancer groups (raw *p*-value < 0.05) with and average effect size of 1.37-fold changes, ranging from 0.59-3.03-fold. The levels of 133 metabolites were higher in ERneg tumors and 31 metabolites were lower in ERneg tumors (Figure 2C, Table S4).

**Figure 2.**
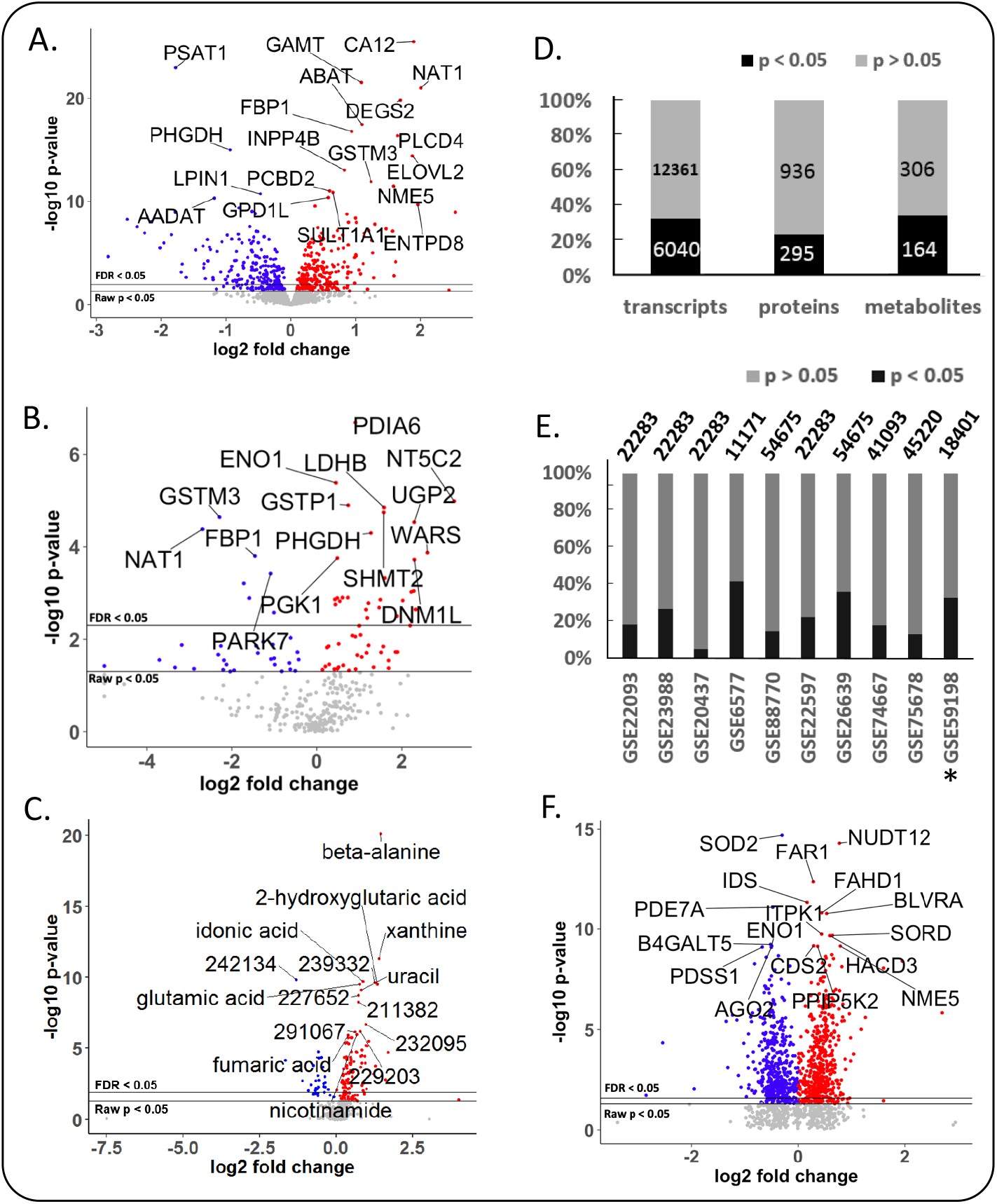
Differential expression of metabolic genes, metabolic enzymes and metabolites in breast cancer, using MetaCancer and GEO studies. Blue: lower in ERneg tumors, Red: higher in ERneg tumors, Grey: no significant change. *Left panel:* MetaCancer multi-omics expression study. (A) transcriptomics data. (B) metabolic enzymes, (C) metabolites. *Right panel:* (D) Overview of MetaCancer transcriptomics, proteomics and metabolomics data. (E) Percentage of significant transcripts across ten GEO studies with the total number of transcripts detected on each bar; * indicates the MetaCancer study. (F) Differential expression of metabolic gene transcripts (raw *p*-value < 0.05) found in at least 7 GEO studies. KS *p*-values calculated from raw *p*-values across ten GEO studies.

The proteomics MetaCancer data consisted of 125 formalin-fixed paraffin embedded (FFPE) samples using 96 ERpos tumors and 29 ERneg tumors. FFPE samples were not available for all 251 patients. ER status for one sample was not known. We could not use the exact same fresh frozen tumors as used for the metabolomics analysis because of limited sample availability. Conversely, FFPE samples are not useful for metabolomics analysis due to the FFPE fixation process. Out of total 1500 detected proteins, 1231 were found in at least six samples in either ERpos or ERneg groups and were kept in the dataset for statistics analysis (Table S5). A total 295 proteins were found to be significantly different regulated (raw *p*-value < 0.05) using the Mann-Whitney U test. The levels of 97 proteins were lower in ERneg tumors and 198 proteins were higher in ERneg tumors (Figure 2B, Table S5).

The transcriptomics MetaCancer dataset was downloaded from the gene expression omnibus (GEO) database with accession number GEO59198. The data set included 18401 genes determined in 122 ERpos and 32 ERneg breast tumors (Table 1). A total of 6040 genes (33%) were significantly different (raw *p*-value < 0.05) between ERpos and ERneg groups using the GEO2R utility (27008011). Among those genes, 3103 were under-expressed and 2937 were over-expressed in the ERneg tumors (Table 1, Table S6). We have then utilized the Expasy database to obtain enzyme commission (EC) annotations for all human genes from NCBI. We selected the EC numbers that chemically transform small-molecules by excluding enzymes that use proteins or genes as substrates, yielding a total of 1549 human genes coding for metabolic enzymes (Table S7). 540 metabolic genes (35%) were found to be significantly different (raw *p-* value < 0.05) in ERneg tumors compared to ERpos tumors in the MetaCancer cohort (Figure 2A, Table S7 and Figure S1).

##### Breast tumor public gene expression datasets

We collected eight additional gene expression datasets for which the ER status was known. The eight additional GEO studies consisted of total 540 for ERpos and 299 ERneg breast cancer samples, ranging between 32-138 samples for ERpos tumors and 10-88 ERneg tumors (Table 1). On average, we found 23% of all transcripts to be significantly different expressed between ERpos and ERneg, ranging from 5-42% (Figure 2E). Similar to the MetaCancer study, 47% of all significant genes were over-expressed, 53% were under-expressed. A meta-analysis by comparing the raw *p*-values distribution of each gene with across all datasets with a null-distribution using the Kolmogrov-Smirnov test (KS) yielded a total of 929 metabolic genes which were significant (raw *p*-value <0.05) (Figure 2F, Table S7, Figure S1). Estrogen receptor 1 (ESR1) gene expression was lower in ERneg tumors across all gene expression datasets (Table S6).

#### Integrated pathway analysis and visualization

Next, we investigated whether these different omics data can be grouped by functional relationships. Here, we focused on integrated analysis of metabolic pathways. We used statistical over-representation as generic tool to summarize and rank the differential regulation of genes, proteins and metabolites by mapping to the Reactome database ^12^. We used Reactome as reference database because it uses GO terms and because it provides more comprehensive mapping of genes to metabolic pathways than KEGG pathways, BioCyc or HMDB. We then confined the Reactome analysis to significantly regulated molecular markers, yielding a joint list of 542 genes, 104 proteins and 56 metabolites. Notably, 20 metabolites, 7 proteins and 45 genes could not be mapped to any pathway set and therefore had to be discarded from further analysis. The lists of remaining molecular markers were summarized by Reactome overrepresentation analysis into 886 pathways (Table S9) of which 88 (~10%) were found to be significantly enriched (Figure 3). Interestingly, for most pathways, the over-representation analysis was based on all three omics levels, while 15 pathways were supported by only protein and gene lists and 8 pathways were based solely on gene set enrichments. This finding is based on the larger size of gene lists compared to proteins or metabolites, in addition to bias in proteomics and metabolomics data acquisitions. For example, the metabolomics platform used here did not cover biosynthesis of complex lipids, while our FFPE-based proteomics method failed to observed low-abundant proteins. We did not use lipidomics analysis here because most lipids are not annotated by specific enzymes in Reactome. Consequently, only gene lists supported the finding of dysregulation of conjugation of amino acids or carboxylic acids (xenobiotic metabolism, Figure 3, branch 5a) and only gene-or gene/protein lists supported the finding of dysregulation of complex lipids (Figure 3, branch 7a). Importantly, Reactome pathway mapping highlighted various biological pathways involved in amino acids (Figure 3, branch 9), carbohydrates (Figure 3, branch 1), nucleotides (Figure 3, branch 3), fatty acyl metabolism (Figure 3, branch 7b), and mitochondrial oxidation (TCA cycle, Figure 3, branch 2) (Table S9). Key specific pathways are within these branches were glycolysis, pentose phosphate pathway, TCA cycle, nucleotide salvage, glutathione conjugation, steroids metabolism, fatty acyl-CoA biosynthesis, serine biosynthesis and metabolism of aromatic amino acids. These pathways were also found to be significantly associated with ERneg phenotype when genes, protein and metabolite lists were analyzed separately (Table S10-S13). Several branches indicate the importance of lipids, nucleotides and amino acids for sustaining tumor growth and cell division in the more aggressive ERneg tumors. Other branches such as carbohydrates and TCA metabolism can be summarized in altered utilization of energy sources, amino acid conjugation for neutralizing xenobiotic including anti-tumor drugs, cholesterol biosynthesis regulation by SREBP and coenzyme biosynthesis to support fatty acid production.

**Figure 3:**
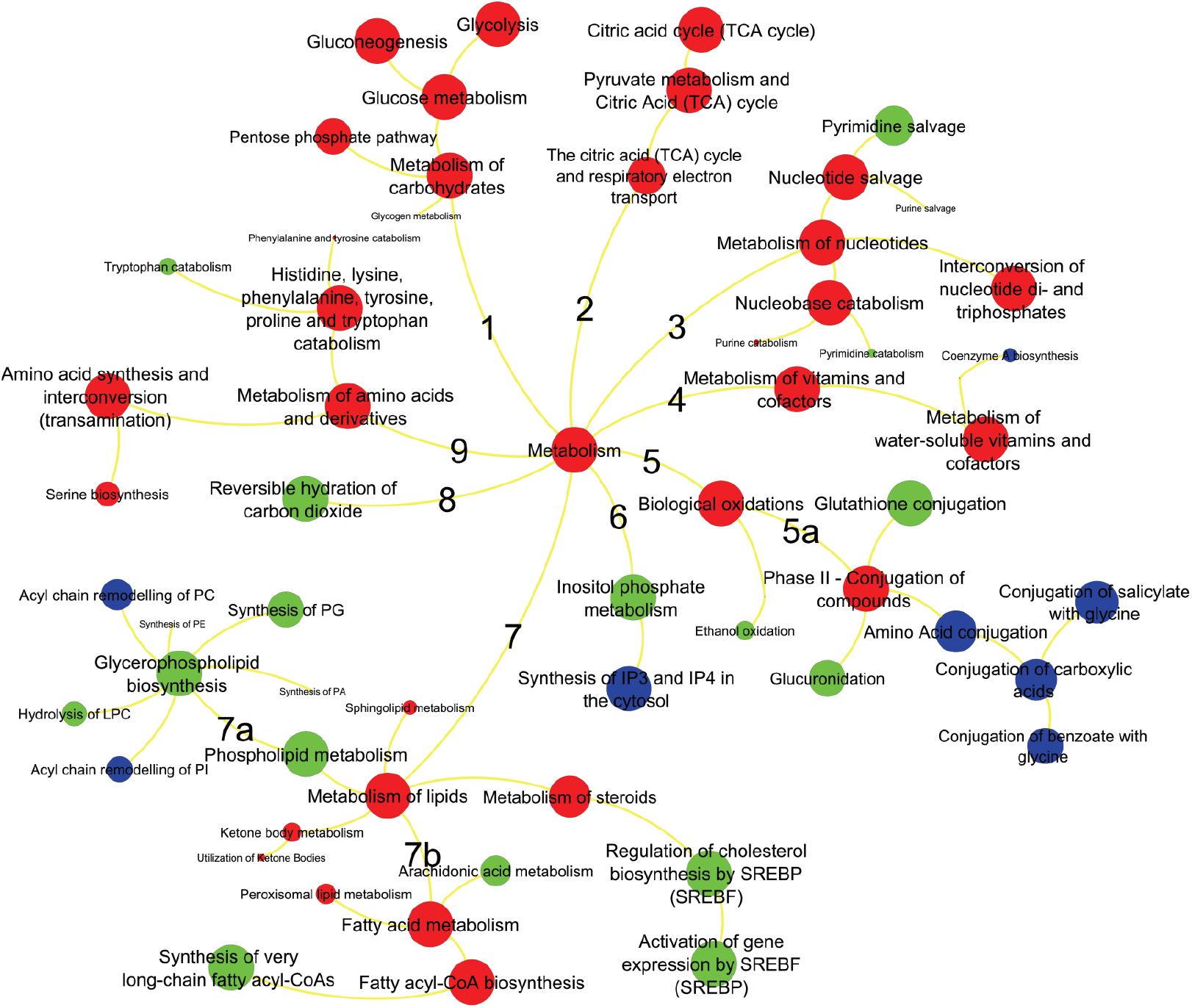
Metabolic pathways significantly enriched in ERneg breast tumors. Key pathways branches are labeled by numbers. Larger node sizes indicate increased significance of pathway dysregulation using the Reactome database tools (29145629, 28249561). Red colors represent pathways supported by gene, protein and metabolites data, green colors are pathways supported by only proteins and genes, blue pathways are based on gene data only.

Next, we integrated differentially regulated genes, proteins and metabolites into network maps (Figure 4) using their chemical and biochemical relationships ^3^. The networks complemented the pathway over-representation analysis by visualizing individual entities and their relationships (also see Figure S2, Table S2). For example, an increase in nucleotide metabolism is substantiated by altered levels of 10 metabolites, 19 metabolic genes and six enzymes including increased levels of 5NTC, NAMPT, BPNT1, PNPH and KCRU (Figures 4, Table S5, S6), in addition to decreased levels of 15 genes (Table S6). Similarly, glycolysis and TCA cycle metabolism (Figure 4) was altered by increased levels of 7 metabolites, 33 genes and 13 proteins, including G6PD, PGM1, MAOM, IDHP, CISY and lower levels of F16P2 (Table S5). The combined action of these genes and proteins supports metabolic reprogramming towards the pentose phosphate pathway (PPP) and TCA anaplerosis. NADPH metabolism (Figure 4) was activated via the PPP to provide more reducing potential for fatty acid synthesis and increased levels of pentose sugars for nucleotide biosynthesis (Table S2). Higher levels of GLYM, P5CS, P5CR1, P4HA1 and PRDX4 indicated increased collagen remodeling, a pathway that was missed by Reactome analysis. To sequester xenobiotics (Figure 4), ERneg tumors preferred conjugation of amino acids by cytochrome P450 genes instead of using glutathione transferases. Not all metabolic dysregulations are represented in Figure 4. For example, DHCR7, involved in cholesterol biosynthesis, was higher in the ERneg tumors (Table S6).

**Figure 4.**
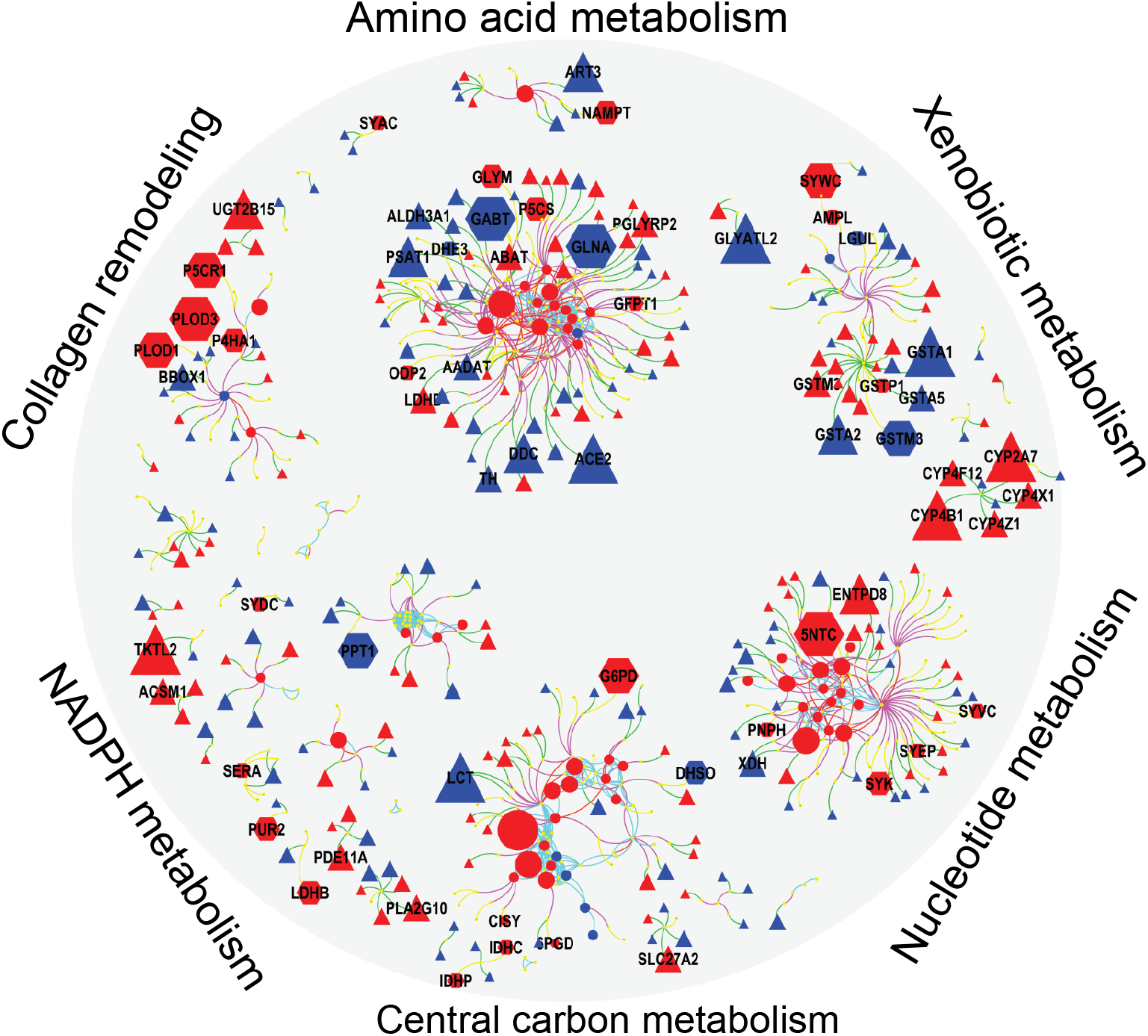
Integrated visualization of genes (Δ), protein 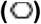 and metabolites (O) by mapping their biochemical and chemical relationships. Edges: Green – gene to enzyme, yellow – protein to enzyme, pink – compound to enzyme, red – compound to compound (biochemical KEGG Rpair), aqua – compound to compound (Tanimoto chemical similarity >0.7). Node colors: blue – lower and red – higher in ERneg breast tumors. Node size reflect the fold changes. Clusters detected by Glay community detection algorithm (29109515).

### Identification of metabolic genes and proteins that were under-studied in breast cancer

Many of the differentially regulated proteins or genes in the network maps (Figure 4) have been studied before. Next, we asked which of these genes and proteins could serve as novel therapeutic targets? To this end, we searched 30 million PubMed abstracts using key-word text-mining to rank all differentially regulated metabolic genes by their publication frequencies in breast tumor biology (see Methods, Table S7). The resulting Table 2 gives the differential expression for the top-50 most significant genes from the MetaCancer and the GEO meta-analysis study in relation to the publication record. This comparison yielded a range of metabolic genes that are currently under-studied with less than five papers. Interestingly, over 2/3 of the 50-most significant metabolic genes are shown here to be severely understudied. Several of the understudied genes are directly involved in critical metabolic pathways of tumor etiology, including glucose metabolism (SORD), lipid biosynthesis (FAR1), cAMP signaling pathway (PDE7A, ADCY6), and glutamate anaplerosis (ABAT, GLUD1). We pose that these under-studied genes should be further investigated in their role in breast tumor aggressiveness. To check the clinical significance of these genes, we have used KMPlot ^19^ tool that estimates the relapse free survival (RFS) using a standardized cohort of breast cancer gene expression studies. We have found that 29 genes were positive associated with RFS and 11 genes negatively associated with RFS in a meta-cohort of 1,809 patients (Supplementary Table S15).

**Table 2.** Prioritization of metabolic candidate genes using omics and text mining. Top 50 most significant genes in the GEO meta-analysis for ten studies and their significance in the MetaCancer proteomics and transcriptomics datasets (raw p-values). Note: - indicates proteins not found in the proteomics dataset

| Gene Symbol | Gene Description | # papers | Meta-Analysis<br>*p*-value | MetaCancer |  |
| --- | --- | --- | --- | --- | --- |
|  |  |  |  | Gene<br>*p*-value | Protein<br>*p*-value |
| SOD2 | superoxide dismutase 2 | 84 | 2.00E-15 | 2.30E-05 | 2.93E-02 |
| NUDT12 | nudix hydrolase 12 | 0 | 5.00E-15 | 7.90E-05 | - |
| FAR1 | fatty acyl-CoA reductase 1 | 0 | 4.50E-13 | 1.80E-03 | - |
| IDS | iduronate 2-sulfatase | 27 | 4.80E-12 | 5.60E-03 | - |
| PDE7A | phosphodiesterase 7A | 0 | 8.20E-12 | 9.20E-10 | - |
| FAHD1 | fumarylacetoacetate hydrolase domain containing 1 | 0 | 1.60E-11 | 2.10E-04 | - |
| BLVRA | biliverdin reductase A | 0 | 1.70E-11 | 4.30E-02 | 1.10E-01 |
| ITPK1 | inositol-tetrakisphosphate 1-kinase | 0 | 1.80E-10 | 2.50E-06 | - |
| SORD | sorbitol dehydrogenase | 0 | 2.10E-10 | 5.90E-06 | 8.14E-02 |
| HACD3 | 3-hydroxyacyl-CoA dehydratase 3 | 0 | 2.20E-10 | 2.40E-02 | - |
| B4GALT5 | beta-1,4-galactosyltransferase 5 | 0 | 5.90E-10 | 4.20E-05 | - |
| CDS2 | CDP-diacylglycerol synthase 2 | 0 | 6.90E-10 | 1.40E-03 | - |
| ENO1 | enolase 1 | 16 | 6.90E-10 | 3.10E-04 | 3.72E-05 |
| NME5 | NME/NM23 family member 5 | 2 | 7.10E-10 | 3.30E-12 | - |
| PPIP5K2 | diphosphoinositol pentakisphosphate kinase 2 | 0 | 7.30E-10 | 9.70E-03 | - |
| PDSS1 | decaprenyl diphosphate synthase subunit 1 | 0 | 7.90E-10 | 4.30E-07 | - |
| AGO2 | argonaute 2, RISC catalytic component | 24 | 8.10E-10 | 9.80E-05 | - |
| ENPP1 | ectonucleotide pyrophosphatase/phosphodiesterase 1 | 9 | 1.90E-09 | 1.90E-04 | - |
| UGP2 | UDP-glucose pyrophosphorylase 2 | 0 | 2.10E-09 | 1.90E-04 | 1.01E-04 |
| COX6C | cytochrome c oxidase subunit 6C | 3 | 2.30E-09 | 1.60E-05 | - |
| GSTA1 | glutathione S-transferase alpha 1 | 24 | 2.60E-09 | 1.10E-07 | 4.71E-01 |
| CSAD | cysteine sulfinic acid decarboxylase | 1 | 2.80E-09 | 3.80E-07 | - |
| GAMT | guanidinoacetate N-methyltransferase | 0 | 3.00E-09 | 2.90E-22 | - |
| CA12 | carbonic anhydrase 12 | 18 | 4.00E-09 | 3.30E-26 | - |
| GLUD1 | glutamate dehydrogenase 1 | 1 | 4.90E-09 | 1.10E-04 | 1.16E-01 |
| LIPG | lipase G, endothelial type | 3 | 5.40E-09 | 6.70E-06 | - |
| SPR | sepiapterin reductase | 88 | 6.50E-09 | 3.90E-06 | 1.20E-01 |
| ME1 | malic enzyme 1 | 6 | 7.00E-09 | 1.60E-04 | 5.24E-02 |
| FBP1 | fructose-bisphosphatase 1 | 13 | 7.50E-09 | 1.60E-17 | 3.51E-03 |
| ABAT | 4-aminobutyrate aminotransferase | 4 | 8.90E-09 | 3.50E-18 | 1.24E-01 |
| PLCH1 | phospholipase C eta 1 | 0 | 1.10E-08 | 1.70E-07 | - |
| PIGH | phosphatidylinositol glycan anchor biosynthesis class H | 2 | 1.20E-08 | 4.90E-05 | - |
| PDXK | pyridoxal kinase | 1 | 2.20E-08 | 2.80E-02 | 1.39E-01 |
| ASAH1 | N-acylsphingosine amidohydrolase 1 | 8 | 2.40E-08 | 7.40E-05 | 2.70E-02 |
| MAN2B2 | mannosidase alpha class 2B member 2 | 0 | 2.40E-08 | 2.30E-03 | - |
| AHCYL1 | adenosylhomocysteinase like 1 | 0 | 3.00E-08 | 3.70E-02 | - |
| COX4I1 | cytochrome c oxidase subunit 4I1 | 2 | 3.10E-08 | 1.30E-01 | - |
| SEPHS1 | selenophosphate synthetase 1 | 0 | 3.40E-08 | 2.00E-03 | 8.86E-01 |
| ATP5S | ATP synthase, H+ transporting, mito-chondrial F<sub>0</sub> complex subunits (factor B) | 0 | 3.70E-08 | 1.20E-03 | - |
| ADCY6 | adenylate cyclase 6 | 1 | 4.20E-08 | 1.90E-05 | - |
| LCMT2 | leucine carboxyl methyltransferase 2 | 0 | 4.60E-08 | 3.30E-08 | - |
| EXT1 | exostosin glycosyltransferase 1 | 10 | 5.20E-08 | 1.70E-01 | - |
| COX6A1 | cytochrome c oxidase subunit 6A1 | 1 | 5.50E-08 | 1.10E-01 | - |
| MTAP | methylthioadenosine phosphorylase | 9 | 6.40E-08 | 4.80E-02 | 4.50E-02 |
| NT5DC2 | 5'-nucleotidase domain containing 2 | 0 | 6.70E-08 | 8.00E-08 | - |
| AK4 | adenylate kinase 4 | 0 | 7.10E-08 | 6.90E-06 | - |
| LPIN1 | lipin 1 | 3 | 8.10E-08 | 1.80E-11 | - |
| GLS | glutaminase | 50 | 9.20E-08 | 3.30E-01 | - |
| RAB6B | RAB6B, member RAS oncogene family | 0 | 9.48E-08 | 8.86E-05 | - |
| PLA2G16 | phospholipase A2 group XVI | 1 | 1.03E-07 | 3.64E-08 | - |

In the context of finding novel, drugable targets for ERneg breast tumors, we then analyzed specifically the proteomics dataset with respect to finding the upregulated proteins that are supported by the GEO meta-analysis, irrespective of the protein function or involvement in metabolism. From a total of 1231 proteins, 295 proteins (24%) were differentially regulated in MetaCancer with a raw *p*-value <0.05. 39 of these proteins were also found to be significantly different in the GEO meta-analysis, of which 21 were up-regulated in ERneg tumors (Figure 5A). Importantly, 10 of these proteins were found to be severely under-studied using our PubMed text mining approach (Figure 5B, Table S14), including CALML5, ACTR3, PADI2 and PFKP. Among these 10 under-studied proteins, four proteins were encoded by metabolic genes, including PFKP, GART, PLOD1 and ASS1. Although PFKP is a key regulatory enzyme in glycolysis and gluconeogeneis pathways, only four PubMed abstracts were found targeting this gene in breast cancer research. We propose to remove bias in study designs to not only study heavily published genes such as CD44 and GSTP1 but also relevant new targets such as PFKP, GART, PLOD1 and ASS1.

**Figure 5.**
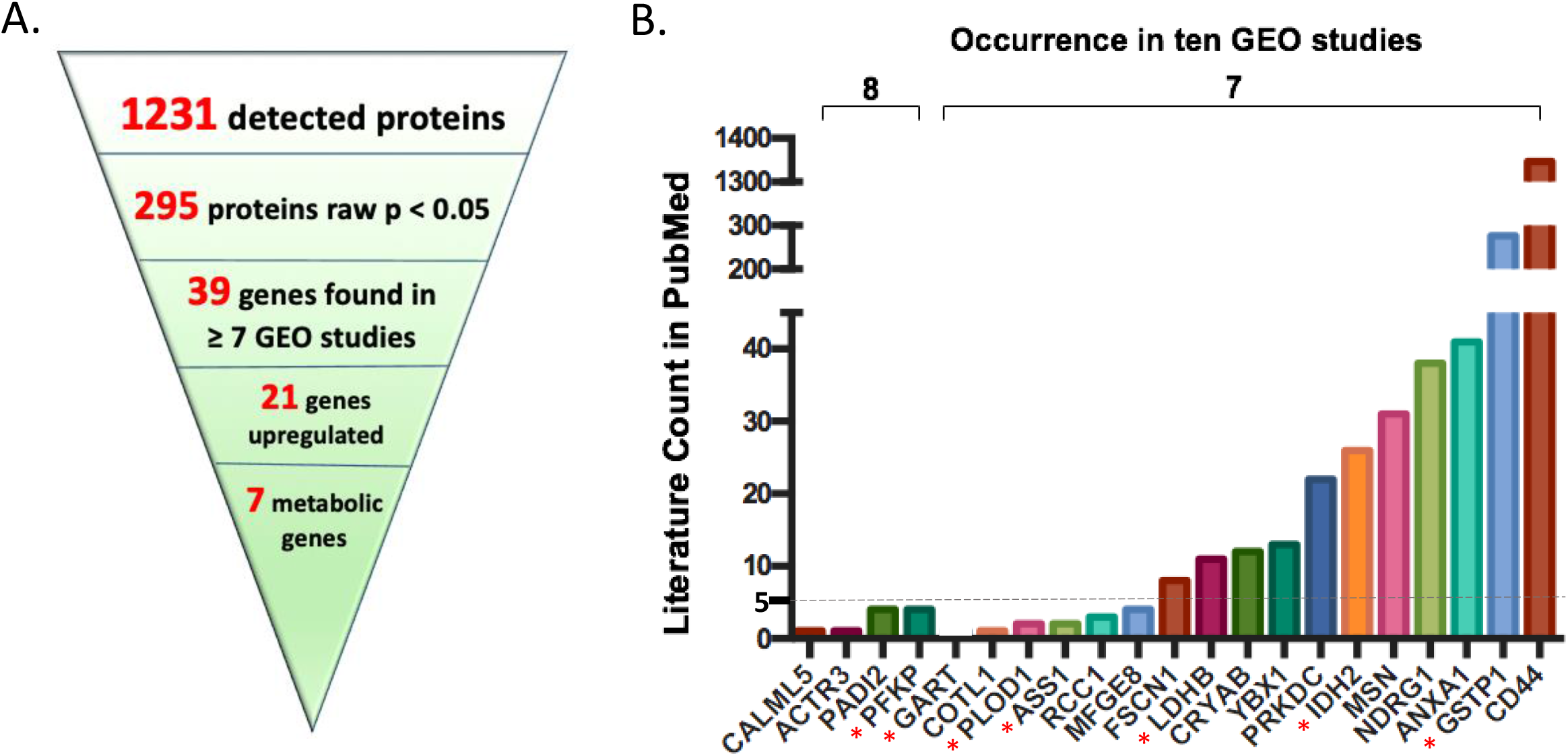
Key upregulated protein encoding genes most often found in ten GEO omnibus studies. 21 key upregulated protein encoding genes were found when liking proteomics dataset with ten GEO omnibus studies (A); PubMed literature counts for most often found genes in ten GEO omnibus studies. * indicates proteins encoded by metabolic genes. Abbreviation: CALML5: Calmodulin-like protein 5; ACTR3: Actin-related protein 3; PADI2: Protein-arginine deiminase type-2; PFKP: Phosphofructokinase, platelet; GART: Trifunctional purine biosynthetic protein adenosine-3; COTL1: Coactosin-like protein; PLOD1: Procollagen-lysine,2-oxoglutarate 5-dioxygenase 1; ASS1: Argininosuccinate synthase; RCC1: Regulator of chromosome condensation; MFGE8: Lactadherin; FSCN1: Fascin; LDHB: L-lactate dehydrogenase B chain; CRYAB: Alpha-crystallin B chain; YBX1: Nuclease-sensitive element-binding protein 1; PRKDC: DNA-dependent protein kinase catalytic subunit; IDH2: Isocitrate dehydrogenase [NADP], mitochondrial; MSN: Moesin; NDRG1: Protein NDRG1; ANXA1: Annexin A1; GSTP1: Glutathione S-transferase P; CD44: CD44 antigen.

**Figure 6.**
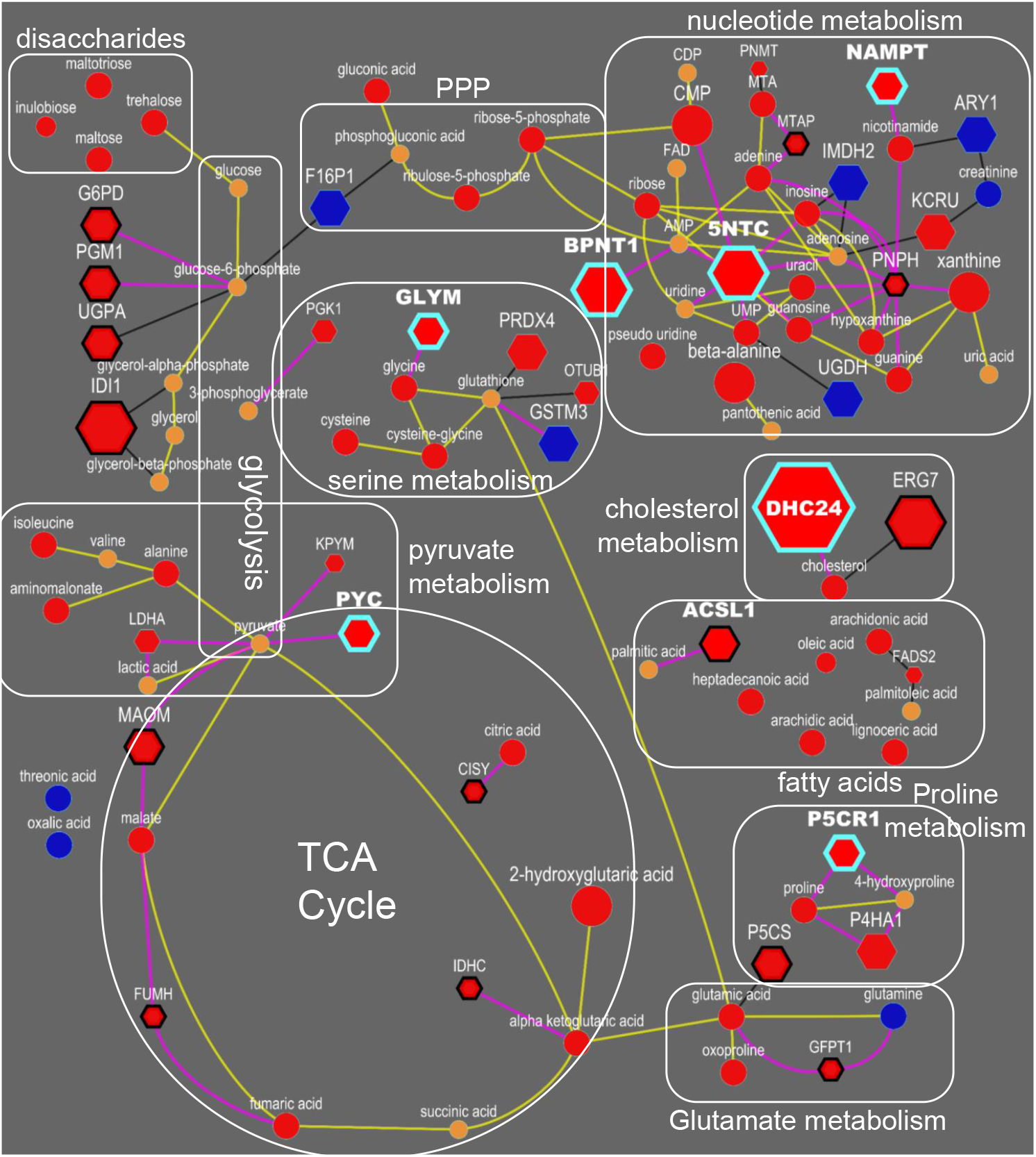
Integrated biochemical pathway visualization for metabolite and proteins datasets. Top-ranked drug targets are highlighted with blue-border. Spheres are metabolites, and hexagons are enzymes. Yellow lines reflect KEGG RPAIR links whereas pink lines reflect reactant-enzyme links. Blue and red color represent significant (p <0.05, ANOVA) decrease and increase in the levels in ER-negative in comparison to ER-positive tumors. Orange color reflects not significant changes in the levels. Size of the nodes reflects fold changes. ER-negative tumors rely more on TCA anaplerosis, anabolic glycolysis, de-novo biosynthesis of amino acids and nucleotide salvage.

## Discussion

Cancer metabolism is radically different from non-malignant cells. Breast cancers can be grouped into different subtypes by presence of receptor genes, stages or grades. We here focused on comparing ERneg and ERpos classes of breast cancer but did not extend this analysis to further subtypes such as triple negative tumors using HER2 and PR status. When designing the study, we queried the GEO database for finding a larger number of studies that had ER-status reported to ensure to have a good base for meta-analysis of significantly expressed metabolic genes. From the nine studies (plus MetaCancer) we downloaded from GEO, only 5 had also information on HER2 status, reducing the power of gene expression meta-analysis. Secondly, several of those studies had already small sample sizes, and if HER2 status was added, the meta-analysis would further lose statistical power. Similarly, our proteomics data set consisted of only 14 triple-negative tumors, again compromising statistical power for finding differences in metabolic genes.

Therefore, we focused on comparing ERneg and ERpos tumors for which we previously showed stark differences between in central metabolism, altered ratios of glutamine/glutamate (glutaminolysis) and beta-alanine accumulation^6^; yet, an integrated analysis with prioritization and a focus on candidate pathways and druggable targets was missing. Despite the established importance of metabolism as hallmark of cancer, few other studies focused on the interplay of genetic mutations and gene expression and actual metabolic phenotypes. Overall, we have found that ERneg tumors show highly active biosynthesis of amino acids, lipids and nucleotides and utilize substrate recycling as well as xenobiotic metabolism to support tumor growth. We consciously constrained our approach to metabolic genes and gene products and here provide the first integrated multi-omic analysis of ERneg versus ERpos tumors. We used solid statistical footing to focus on interpretations of metabolic differences to find novel targets and give the first example how in-depth text mining can be used to prioritize gene targets that have been largely ignored in the literature. Yet, despite clear evidence from the data, not all metabolic enzymes may be suitable as drug targets because they support critical metabolic pathways in cells throughout the body. Hence, strong inhibition of such enzymes might lead to severe therapeutic side-effects. Instead, knowledge about specific pathways might also be useful for targeting regulatory and signaling pathways in metabolism that may have less toxicity in normal cells. Our study results presented here support the idea that specific metabolic genes have gained less attention than others, even if they act in the same pathway. For example, our analysis showed that glutaminase is frequently reported in PubMed abstract in reference to breast cancer, whereas the classic mitochondrial glutamate oxidation enzyme, GLUD1, has only been once reported in this context. Specifically, in relation to new research on targeted drug delivery may render high-flux metabolic enzymes equally important for new therapy options as immunotherapy or classic oncology chemotherapies.

### Nucleotide salvage pathway

Reactome pathway analysis of our data highlighted the nucleotide salvage pathway as specifically important in ERneg tumors (Figure 3, branch 3, Table S9). 5NTC (Cytosolic purine 5’-nucleotidase), a key enzyme involved in nucleotide salvage, was up-regulated in ER-negative tumors. Five purine metabolites were elevated in ER-negative tumors, including adenine, guanosine, guanine, xanthine and hypoxanthine (Figure 5). Meanwhile, elevated levels of beta-alanine were observed in ER-negative tumors, an intermediate of the pyrimidine salvage pathway, along with the concurrent increases of uracil, pseudo-uridine, UMP and CMP. Concurrently, 5-deoxy-methylthioadenosine (MTA), a further purine salvage metabolite, was also observed at higher levels in ER-negative tumors (Table S4). Aggressive tumors bypass autophagy and apoptosis and hence, require more re-use of nucleotides for cell survival and cell division. This process may therefore contribute to the cancer phenotype of cell survival. In addition, cancer cells also activate *de novo* nucleotide biosynthesis. There are a range of drugs targeting these metabolic pathways in chronic lymphocytic leukaemia, lung cancer and pancreatic cancer ^28^, but these drugs have not yet been repurposed to be tested against ER-negative tumors. Our analysis motivates and supports the initiation of such clinical trials of these drugs for the management of ERneg breast tumors.

### Microenvironment remodeling

Metabolites involved in collagen biosynthesis as well as collagen remodeling enzymes were enriched in ER-negative tumors. Breast tumor cells increasingly rely on *de novo* biosynthesis of proline for collagen metabolism ^37^. Accordingly, we found increased levels of proline and trans-4-hydroxyproline as well as two enzymes involved in *de novo* biosynthesis of proline in ERneg tumors, pyrroline-5-carboxylate reductase (PYCR) and P5C-synthase (ALDH18A1) (Table S5). PYCR converts pyrroline-5-carboxylate (P5C) to proline and ALDH18A1 produce P5C from glutamate.

Extracellular matrix (ECM) is the most abundant component in the tumor microenvironment and it has been associated with breast cancer progression and metastatic spread^35^. As a scaffold of tumor microenvironment, collagen changes in the microenvironment regulate ECM remodeling, release signals and trigger a cascade of biological events, promote tumor invasion and migration^14^. ECM proteins and ECM mediated signaling pathways may be promising drug targets for breast cancer ^23^. Along with the change of collagen genes, the expression levels some chaperone and co-chaperone proteins, such as HSP90AA1, HSP90AB1, HSP90B1 and CDC37, were higher in ERneg tumors, as well as YWHAQ, YWHAE (Supplemental Table S9), which involved in PI3K-AKT signaling pathway ^40^.

### Carbohydrate metabolism

The strongest effect supported by three omics levels (metabolites, proteins and genes) was found for an increased PPP activity. PPP intermediates, ribose-5-phosphate and ribulose-5-phosphate, were increased in ERneg tumors, caused by an increase in the abundance of the key oxidation enzymes, glucose-6-phosphate dehydrogenase (G6PD) and 6-phosphogluconate dehydrogenase (PGD) and their encoding genes. Levels of transketolase, a key enzyme in the non-oxidative branch of the pentose phosphate pathway, and its encoding gene TKT, were also found increased in ERneg tumors. Further, gene expressions of phosphoglucomutase 1 (PGM1), ribose-5-phosphate isomerase (RPIA) and deoxyribose-phosphate aldolase (DERA) were also upregulated in ERneg tumors (Table S5, 7). The pentose phosphate pathway delivers both NADPH and pentose phosphates required for cell division and tumor proliferation^26^. The activation of this pathway supports tumor growth known for ERneg tumors. In comparison to the clear differential regulation of the PPP, glycolytic enzymes showed less consistent regulation of genes, proteins and metabolites when comparing ERneg to ERpos tumors.

Apart from metabolic changes in PPP, higher levels of three uncommon carbohydrates were observed, including a two-fold increase in trehalose, maltose and maltotriose levels in ERneg tumors (Table S4 and Figure 5). These oligosaccharides might be co-imported from blood along with the well-known glucose uptake in tumors. This is the first report on these compounds in relation to breast tumor metabolism; the maltose-degrading enzyme (GAA, LYAG) in the MetaCancer study was found down-regulated (Table S5) while the corresponding gene was found significantly down-regulated, consistent with other studies in the GEO meta-analysis (Table S7). This significance in differential regulation of oligosaccharide metabolism in ERneg tumors on all three omics levels suggests that further potentially this pathway could also be important for future drug therapies.

### Mitochondrial oxidation

We have observed increased levels of the metabolites nicotinamide and citric acid cycle (TCA) intermediates citrate, alpha-ketoglutarate, malate, fumarate and succinate (Table S4) along with over-expression of citrate synthase (CISY) and Nicotinamide phosphoribosyltransferase (NAMPT) in ERneg breast cancer patients (Table S5). Under hypoxic condition, attenuation of electron transport chain in tumors is expected and NADH production in TCA can imbalance the NAD+/NADH ratio. Targeting the NAD+ salvage pathway is a promising therapeutic option for cancer patients^15^. Aggressive breast tumor shows an overexpression of hypoxia inducible factor 1 alpha (HIF-1)^4^. NAMPT also known as visfatin, the rate-limiting enzyme in the NAD salvage pathway has been targeted by inhibitors such as FK866 ^21^ and CHS-828^34^. This protein has also been reported to be overexpressed in prostate cancer to promote tumor cell survival ^42^ and is required for *de novo* lipogenesis in the tumor cells^5^. Over-expression of NAMPT in breast cancer tissue is associated with poor survival ^30^. Inhibiting NAMPT can also make the ERneg breast cancer sensitive to additional chemotherapies as observed *in vitro* ^2^.

### Poorly studied metabolic genes

Transcriptomics analysis often reveals a long list of significant regulated genes, some of which could be drivers for poor outcomes in subjects with ERneg tumors, while other genes might be regulated downstream as bystanders. A focus on metabolic genes, combined with meta-analysis and multi-omics integration, is proposed here to remove bias in candidate gene selections in the breast cancer research community. Gene expression meta-analysis yielded 34 significant metabolic genes that are underreported with fewer than five PubMed abstracts as targets in breast cancer research, including NUDT12, FAR1, PDE7A, FAHD1, BLVRA, ITPK1, SORD, HACD3, B4GALT5, CDS2, PPIP5K2, PDSS1, UGP2, GAMT, PLCH1, MAN2B2, AHCYL1, SEPHS1, ATP5S, LCMT2, NT5DC2, AK4, CSAD, GLUD1, PDXK, ADCY6, COX6A1, NME5, PIGH, COX4I1, COX6C, LIPG, LPIN1 and ABAT (Table 3). In addition, the combination of proteomics and gene meta-analysis results revealed that 4 upregulated proteins encoded by metabolic genes, PFKP, GART, PLOD1 and ASS1, were under-studied (Figure 5B).

For example, higher expression levels of gene encoding ATP-dependent 6-phosphofructokinase, platelet type (PFKP) were found to be statistically up-regulated in ERneg tumors in eight out of ten GEO studies and specifically, was also found up-regulated in both the proteomics and transcriptomics dataset of the MetaCancer cohort (Figure 5B). However, PubMed text mining showed there were only four publications focused on PFKP in reference to breast cancer. PFKP is a critical rate limiting enzyme in glycolysis, phosphorylating fructose-6-phosphate to fructose-1,6-bisphosphate and determining the rate of glycolytic flux versus flux into the pentose phosphate pathway. Two further 6-phosphofructokinase isomers are existing in humans, PFKM (muscle type) and PFKL (liver type)^41^. PFKM was not detected in our proteomics dataset and not significantly different in transcriptomics dataset (raw *p* = 0.08), while the gene expression of PFKL was upregulated in the MetaCancer ERneg breast tumor transcriptomics dataset (raw *p* < 0.05). Therefore, targeting specific isoforms of phosphofructokinase may be useful as potential target to deprive cancer cells from essential substrates and energy for proliferation while allowing the survival of normal cells^20^.

Similarly, other underreported metabolic genes might serve as new therapeutic targets. Three isoforms of procollagen-lysine, 2-oxoglutarate 5-dioxygenase (PLOD) have been identified. PLOD1 hydroxylates a lysine residue in the alpha-helical or central domain of procollagens; PLOD2 is responsible for lysine hydroxylation in the telopeptide of procollagens whereas the substrate specificity of PLOD3 is unknown^16^. PLOD1 and PLOD2 expression was induced by hypoxia in breast cancer cells ^16^. The expression of PLOD1 was upregulated in lysyl oxidase-like 4 (LOXL4) knockout xenograft tumor tissues and LOX4 knockdown could enhance tumor growth and metastasis through collagen-dependent extracellular matrix changes in TNBC^10^. We therefore pose that PLOD isoforms may be interesting targets to study the relevance of metabolism in the tumor microenvironment.

A third example is Argininosuccinate synthetase 1 (ASS1). ASS1 converts citrulline to arginine and is a key enzyme in arginine biosynthesis that is critical for many functions, including nitric oxide signaling or polyamine biosynthesis, as well as the liver urea cycle. In the MetaCancer cohort, we found both ASS1 transcripts and the enzyme up-regulated in ERneg tumors (Table S5, S7), and the ASS1 substrate citrulline being decreased in the metabolomic results (Table S4). An inhibition should restrict arginine availability. Arginine starvation has been shown to impair mitochondrial respiratory function in ASS1-deficient breast cancer cells^36^, suggesting that arginine starvation therapy such as pegylated recombinant arginine deiminase (ADI-PEG20) could be an option for patients with low ASS1 expression^43^. Dietary arginine restriction has also been shown to reduce tumor growth in a xenograft model of ASS1-deficient breast cancer^9^. We pose that such drugs with undergoing clinical trials could be repurposed for ERneg tumor therapy.

### Conclusions

We here show how multi-omics data can be utilized along with text mining to identify metabolic genes and metabolic pathways that are under-studied in breast tumor research, specifically for ERneg tumors that are known to have poor clinical outcomes. We see this approach as a hypothesis-generating method, prioritizing the multitude of genes that are to be significant in classic transcriptomics studies. We also showed that a meta-analysis of multiple studies refines and strengthens candidate gene selections to study metabolic reprogramming. We propose that such understudied, significant metabolic genes could be used as additional potential therapy targets, especially if genes and proteins were found up-regulated in different breast cancer studies, and if metabolite abundance support protein activities.

## Material and Methods

### MetaCancer cohort details

The study included 276 fresh frozen breast tumor biopsies and 126 FFPE tissues. They were collected for the tissue bank of the European FP7 MetaCancer consortium at the Charité Hospital. The project was approved by the institutional review board of the Charité Hospital (EA1/139/05). Further details about the cohort can be found in our previous report (PMID:24125731). Biopsies were used for transcriptomics and metabolomics analysis and FFPE tissues were used proteomics analysis.

### MetaCancer metabolomics and proteomics data

GC-TOF MS data acquisition of the fresh frozen tumor biopsies tissues was performed as previously published (22823888). FFPE tissues were analyzed using a LTQ mass spectrometer for measuring proteins. Details are given in the Supplementary Text S1.

### Public breast tumor transcriptomics datasets

The MetaCancer transcriptomics dataset was downloaded from the GEO omnibus database using the accession number GSE59198. Nine other gene expression datasets were selected for which estrogen receptor status was available in the GEO omnibus database. Details about these datasets are provided in the Table S1.

### Bioinformatics and statistical analysis

Metabolite enrichment analysis was performed using the ChemRICH tool (29109515). Input file for the ChemRICH analysis is provided in the Table S2. Metabolic network mapping was performed using the MetaMapp software (22591066) and the input file is provided in the Table S3. MetaMapp network was visualized using the Cytoscape software. Metabolites were linked to proteins using the KEGG and Expasy databases. Links were visualized as an integrated network using the Cytoscape network visualization software. Metabolites were linked to enzyme commission (EC) number first using the KEGG database and EC number to protein mapping was obtained from the Expasy database. Pathway over-representation analysis was conducted using the Reactome pathway analysis tool (29145629, 28249561) because it provides the most comprehensive coverage of metabolic pathway maps. Gene symbols and “breast cancer” were searched in the PubMed database to get the count of abstracts for a gene and breast cancer. NCBI eutils web services were used for running the searches for all the metabolic genes. All statistical analyses were conducted using R. Mann-Whitney-Wilcoxon test was performed on both metabolomics and proteomics datasets. GEO2R utility was used to analyze transcriptomics datasets.

Notes: Dinesh Kumar Barupal and Bei Gao contributed equally to this work.

## Availability of data and materials

All the datasets are provided in the supplementary section.

## Author’s contribution

OF, DKB, BG, JB and CD designed the study. CD provided the tumor biopsies. OF, DKB, BSP, BP and BG acquired metabolomics and proteomics datasets and performed the statistical analyses. DKB, BG and OF created the integrated graphs and interpret the statistical results. DKB, BG, OF and CD wrote the manuscript. All authors edited and approved the final manuscript.

## Supplementary Materials

### Supplementary Text

Text S1 : Detailed method description for metabolomics and proteomics analysis.

### Supplementary Figures

Figure S1. Overview of differential gene expression for all and metabolic genes in the ERneg and ERpos tumors.

Figure S2. MetaMapp and ChemRICH analysis of the metabolomics datasets. Upper panel : MetaMapp metabolic network mapping of metabolomics datasets. Each node is metabolite. Read mean higher in ERneg tumors and blue means lower. Size of the node shows fold-change difference. Red lines are known biochemical links and green lines are chemical similarity links. Lower panel: Chemical similarity enrichment analysis (29109515) of the metabolic differences between ERneg and ERpos breast tumors. Point size shows the chemical class size and color shows the overall direction of dysregulated metabolites where red mean mostly upregulated and blue means mostly down-regulated metabolites.

### Supplementary Tables

Table S1 Description of GEO omnibus datasets

Table S2 ChemRICH input

Table S3 MetaMapp input

Table S4 Metabolomics dataset

Table S5 Proteomics dataset

Table S6 Transcritomics analysis results and meta-analysis

Table S7 Metabolic genes

Table S8 Input reactome pathway analysis

Table S9 Reactome results-all entities

Table S10 Reactome results-metabolites

Table S11 Reactome results-proteins

Table S12 Reactome results-transcriptomics

Table S13 Reactome results-meta-analysis

Table S14 Literature count for 21 upregulated proteins

Table S15 Association of top 50 metabolic gene expression with breast cancer patient survival in the http://kmplot.com/database.

## References

1 Auslander N, Yizhak K, Weinstock A, Budhu A, Tang W, Wang XW et al. A joint analysis of transcriptomic and metabolomic data uncovers enhanced enzyme-metabolite coupling in breast cancer. Scientific reports 2016; 6: 29662–29662.

2 Bajrami I, Kigozi A, Van Weverwijk A, Brough R, Frankum J, Lord CJ et al. Synthetic lethality of PARP and NAMPT inhibition in triple-negative breast cancer cells. EMBO molecular medicine 2012; 4: 1087–1096.

3 Barupal DK, Haldiya PK, Wohlgemuth G, Kind T, Kothari SL, Pinkerton KE et al. MetaMapp: mapping and visualizing metabolomic data by integrating information from biochemical pathways and chemical and mass spectral similarity. BMC bioinformatics 2012; 13: 99–99.

4 Bos R, Zhong H, Hanrahan CF, Mommers ECM, Semenza GL, Pinedo HM et al. Levels of HypoxiaInducible Factor-1α During Breast Carcinogenesis. JNCI: Journal of the National Cancer Institute 2001; 93: 309–314.

5 Bowlby SC, Thomas MJ, D’Agostino RB, Jr., Kridel SJ. Nicotinamide phosphoribosyl transferase (Nampt) is required for de novo lipogenesis in tumor cells. PloS one 2012; 7: e40195–e40195.

6 Budczies J, Brockmöller SF, Müller BM, Barupal DK, Richter-Ehrenstein C, Kleine-Tebbe A et al. Comparative metabolomics of estrogen receptor positive and estrogen receptor negative breast cancer: alterations in glutamine and beta-alanine metabolism. Journal of Proteomics 2013; 94: 279–288.

7 Budczies J, Pfitzner BM, Györffy B, Winzer K-J, Radke C, Dietel M et al. Glutamate enrichment as new diagnostic opportunity in breast cancer. International Journal of Cancer 2014; 136: 1619–1628.

8 Budczies J, Denkert C. Tissue-Based Metabolomics to Analyze the Breast Cancer Metabolome. Metabolism in Cancer 2016; 207: 157–175.

9 Cheng CT, Qi Y, Wang YC, Chi KK, Chung Y, Ouyang C et al. Arginine starvation kills tumor cells through aspartate exhaustion and mitochondrial dysfunction. Commun Biol 2018; 1: 178.

10 Choi SK, Kim HS, Jin T, Moon WK. LOXL4 knockdown enhances tumor growth and lung metastasis through collagen-dependent extracellular matrix changes in triple-negative breast cancer. Oncotarget 2017; 8: 11977–11989.

11 de Cremoux P, Valet F, Gentien D, Lehmann-Che J, Scott V, Tran-Perennou C et al. Importance of pre-analytical steps for transcriptome and RT-qPCR analyses in the context of the phase II randomised multicentre trial REMAGUS02 of neoadjuvant chemotherapy in breast cancer patients. BMC cancer 2011; 11: 215–215.

12 Fabregat A, Jupe S, Matthews L, Sidiropoulos K, Gillespie M, Garapati P et al. The Reactome Pathway Knowledgebase. Nucleic acids research 2018; 46: D649–D655.

13 Fahrmann JF, Grapov DD, Wanichthanarak K, DeFelice BC, Salemi MR, Rom WN et al. Integrated Metabolomics and Proteomics Highlight Altered Nicotinamide- and Polyamine Pathways in Lung Adenocarcinoma. Carcinogenesis 2017; 38: 271–280.

14 Fang M, Yuan J, Peng C, Li Y. Collagen as a double-edged sword in tumor progression. Tumour biology : the journal of the International Society for Oncodevelopmental Biology and Medicine 2014; 35: 2871–2882.

15 Fleischer TC, Murphy BR, Flick JS, Terry-Lorenzo RT, Gao Z-H, Davis T et al. Chemical Proteomics Identifies Nampt as the Target of CB30865, An Orphan Cytotoxic Compound. Chemistry & Biology 2010; 17: 659–664.

16 Gilkes DM, Bajpai S, Wong CC, Chaturvedi P, Hubbi ME, Wirtz D et al. Procollagen lysyl hydroxylase 2 is essential for hypoxia-induced breast cancer metastasis. Molecular cancer research : MCR 2013; 11: 456–466.

17 Graham K, de las Morenas A, Tripathi A, King C, Kavanah M, Mendez J et al. Gene expression in histologically normal epithelium from breast cancer patients and from cancer-free prophylactic mastectomy patients shares a similar profile. British journal of cancer 2010; 102: 1284–1293.

18 Gruvberger-Saal SK, Bendahl P-O, Saal LH, Laakso M, Hegardt C, Edén P et al. Estrogen Receptor β Expression Is Associated with Tamoxifen Response in ERα-Negative Breast Carcinoma. Clinical Cancer Research 2007; 13: 1987.

19 Gyorffy B, Lanczky A, Eklund AC, Denkert C, Budczies J, Li Q et al. An online survival analysis tool to rapidly assess the effect of 22,277 genes on breast cancer prognosis using microarray data of 1,809 patients. Breast Cancer Res Treat 2010; 123: 725–731.

20 Hasawi NA, Alkandari MF, Luqmani YA. Phosphofructokinase: A mediator of glycolytic flux in cancer progression. Critical Reviews in Oncology/Hematology 2014; 92: 312–321.

21 Hasmann M, Schemainda I. FK866, a Highly Specific Noncompetitive Inhibitor of Nicotinamide Phosphoribosyltransferase, Represents a Novel Mechanism for Induction of Tumor Cell Apoptosis. Cancer Research 2003; 63: 7436.

22 Horvath A, Pakala SB, Mudvari P, Reddy SDN, Ohshiro K, Casimiro S et al. Novel insights into breast cancer genetic variance through RNA sequencing. Scientific reports 2013; 3: 2256–2256.

23 Insua-Rodríguez J, Oskarsson T. The extracellular matrix in breast cancer. Advanced Drug Delivery Reviews 2016; 97: 41–55.

24 Iwamoto T, Bianchini G, Booser D, Qi Y, Coutant C, Ya-Hui Shiang C et al. Gene Pathways Associated With Prognosis and Chemotherapy Sensitivity in Molecular Subtypes of Breast Cancer. JNCI: Journal of the National Cancer Institute 2011; 103: 264–272.

25 Iwamoto T, Bianchini G, Qi Y, Cristofanilli M, Lucci A, Woodward WA et al. Different gene expressions are associated with the different molecular subtypes of inflammatory breast cancer. Breast cancer research and treatment 2011; 125: 785–795.

26 Jeon S-M, Chandel NS, Hay N. AMPK regulates NADPH homeostasis to promote tumour cell survival during energy stress. Nature 2012; 485: 661–665.

27 Jin M-S, Lee H, Woo J, Choi S, Do MS, Kim K et al. Integrated Multi-Omic Analyses Support Distinguishing Secretory Carcinoma of the Breast from Basal-Like Triple-Negative Breast Cancer. PROTEOMICS – Clinical Applications 2018; 12: 1700125.

28 Jordheim LP, Durantel D, Zoulim F, Dumontet C. Advances in the development of nucleoside and nucleotide analogues for cancer and viral diseases. Nature Reviews Drug Discovery 2013; 12: 447.

29 Lawrence RT, Perez EM, Hernandez D, Miller CP, Haas KM, Irie HY et al. The proteomic landscape of triple-negative breast cancer. Cell Rep 2015; 11: 630–644.

30 Lee Y-C, Yang Y-H, Su J-H, Chang H-L, Hou M-F, Yuan S-SF. High Visfatin Expression in Breast Cancer Tissue Is Associated with Poor Survival. Cancer Epidemiology Biomarkers && Prevention 2011; 20: 1892.

31 Li M, Song Y, Cho N, Chang JM, Koo HR, Yi A et al. An HR-MAS MR metabolomics study on breast tissues obtained with core needle biopsy. PloS one 2011; 6: e25563–e25563.

32 Metzger-Filho O, Michiels S, Bertucci F, Catteau A, Salgado R, Galant C et al. Genomic grade adds prognosOc value in invasive lobular carcinoma†. Annals of Oncology 2013; 24: 377–384.

33 Mishra P, Ambs S. Metabolic Signatures of Human Breast Cancer. Molecular & cellular oncology 2015; 2: e992217.

34 Olesen UH, Christensen MK, Björkling F, Jäättelä M, Jensen PB, Sehested M et al. Anticancer agent CHS-828 inhibits cellular synthesis of NAD. Biochemical and Biophysical Research Communications 2008; 367: 799–804.

35 Oskarsson T. Extracellular matrix components in breast cancer progression and metastasis. The Breast 2013; 22: S66–S72.

36 Qiu F, Chen Y-R, Liu X, Chu C-Y, Shen L-J, Xu J et al. Arginine starvation impairs mitochondrial respiratory function in ASS1-deficient breast cancer cells. Science signaling 2014; 7: ra31–ra31.

37 Richardson AD, Yang C, Osterman A, Smith JW. Central carbon metabolism in the progression of mammary carcinoma. Breast cancer research and treatment 2008; 110: 297–307.

38 Schramm G, Surmann EM, Wiesberg S, Oswald M, Reinelt G, Eils R et al. Analyzing the regulation of metabolic pathways in human breast cancer. BMC Med Genomics 2010; 3: 39.

39 She Q-B, Gruvberger-Saal SK, Maurer M, Chen Y, Jumppanen M, Su T et al. Integrated molecular pathway analysis informs a synergistic combination therapy targeting PTEN/PI3K and EGFR pathways for basal-like breast cancer. BMC cancer 2016; 16: 587–587.

40 Slattery ML, Mullany LE, Sakoda LC, Wolff RK, Stevens JR, Samowitz WS et al. The PI3K/AKT signaling pathway: Associations of miRNAs with dysregulated gene expression in colorectal cancer. Molecular carcinogenesis 2018; 57: 243–261.

41 Sola-Penna M, Da Silva D, Coelho WS, Marinho-Carvalho MM, Zancan P. Regulation of mammalian muscle type 6-phosphofructo-1-kinase and its implication for the control of the metabolism. IUBMB Life 2010; 62: 791–796.

42 Wang B, Hasan MK, Alvarado E, Yuan H, Wu H, Chen WY. NAMPT overexpression in prostate cancer and its contribution to tumor cell survival and stress response. Oncogene 2010; 30: 907.

43 Yeh T-H, Chen Y-R, Chen S-Y, Shen W-C, Ann DK, Zaro JL et al. Selective Intracellular Delivery of Recombinant Arginine Deiminase (ADI) Using pH-Sensitive Cell Penetrating Peptides To Overcome ADI Resistance in Hypoxic Breast Cancer Cells. Molecular pharmaceutics 2016; 13: 262–271.

44 Yoon H, Yoon D, Yun M, Choi JS, Park VY, Kim E-K et al. Metabolomics of Breast Cancer Using High-Resolution Magic Angle Spinning Magnetic Resonance Spectroscopy: Correlations with 18F-FDG Positron Emission Tomography-Computed Tomography, Dynamic Contrast-Enhanced and Diffusion-Weighted Imaging MRI. PloS one 2016; 11: e0159949–e0159949.

