## Supplementary Figure S1 for "Prioritization of metabolic genes as novel therapeutic targets in estrogen-receptor negative breast tumors using multi-omics data and text mining"

### Slide 1
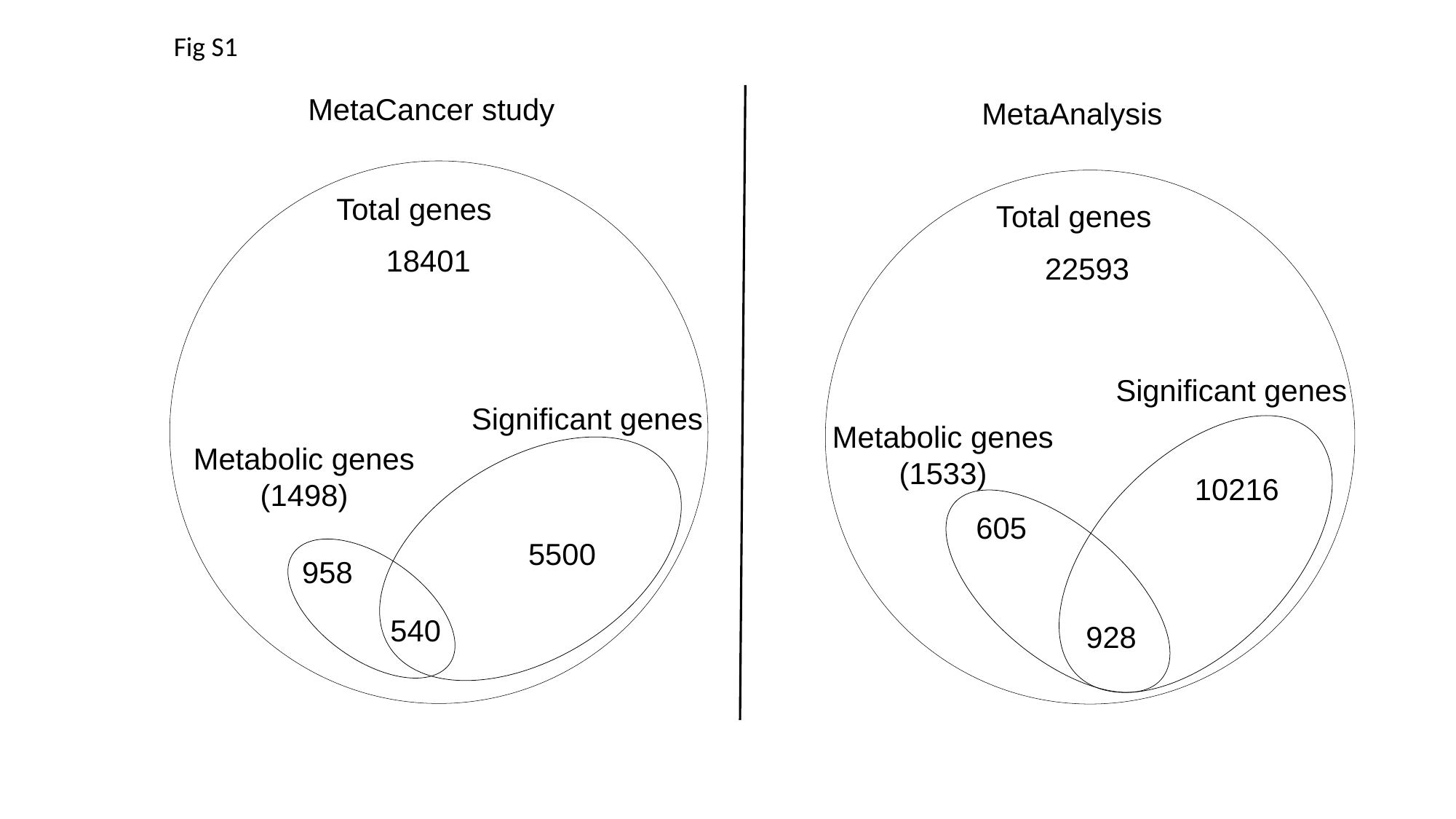

Fig S1
MetaCancer study
MetaAnalysis
Total genes
18401
Significant genes
Metabolic genes
(1498)
5500
958
540
Total genes
22593
Significant genes
Metabolic genes
(1533)
10216
605
928
