## Supplementary figures and images for "Prioritization of metabolic genes as novel therapeutic targets in estrogen-receptor negative breast tumors using multi-omics data and text mining"

### Supplementary Figure S2

#### Slide 1
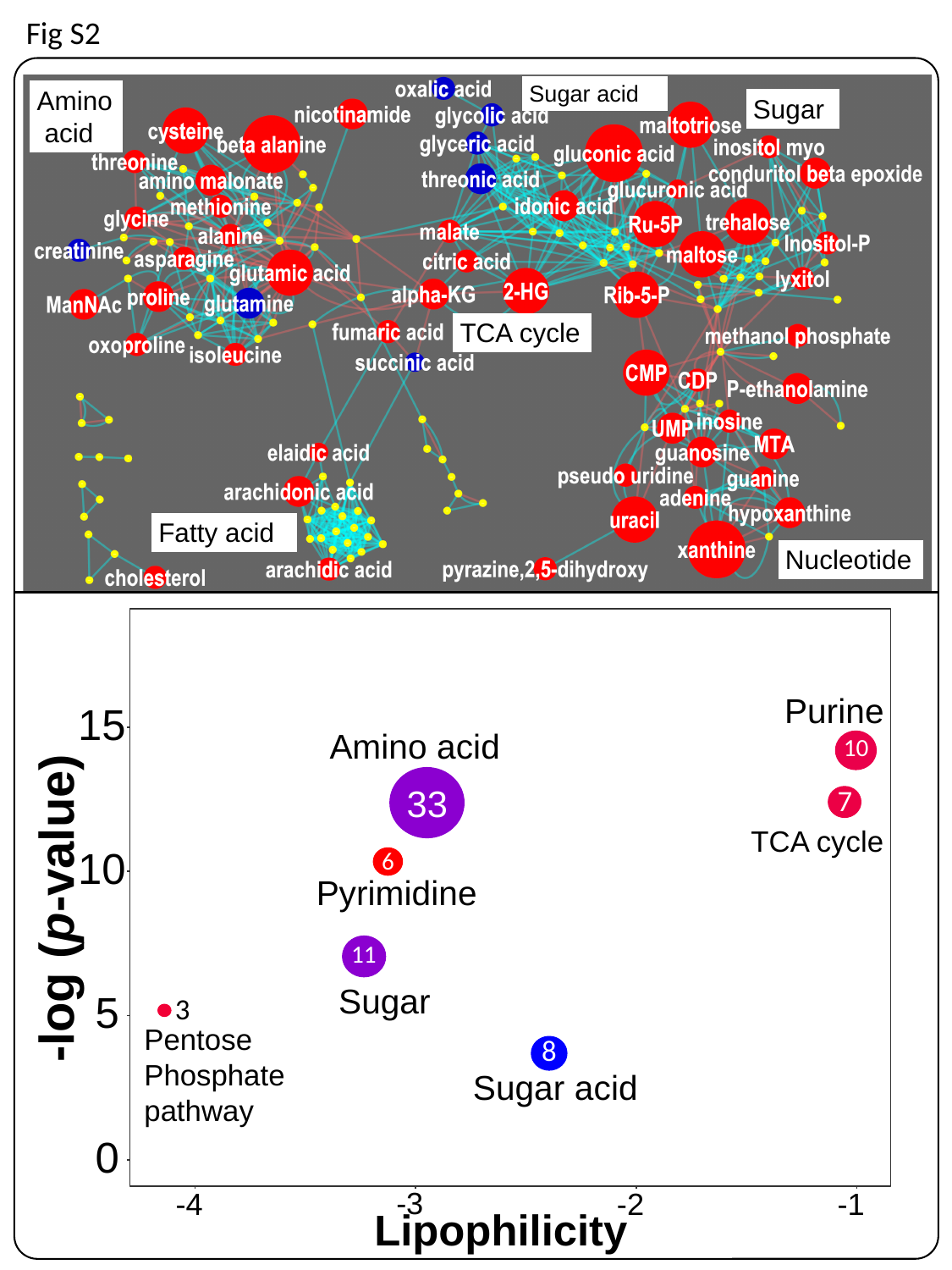

Fig S2
