## Supplementary Text S1 for "Prioritization of metabolic genes as novel therapeutic targets in estrogen-receptor negative breast tumors using multi-omics data and text mining"

Metabolomics method

Briefly, 20 mg frozen breast tissue samples were homogenized and extracted with 1 ml degassed isopropanol/acetonitrile/water (3/3/2) at 4°C for 5 min. Samples were vortexed for 10 s, shaken for 5 min and then centrifuged for 2 min at 14,000 rcf. Two 450 μL supernatant aliquots were transferred to new tubes. One tube was stored as a backup aliquot and another was dried down using CentriVap. The extracts were re-suspended in 50% aqueous acetonitrile to remove most of the complex lipids. After dry evaporation, extracts were derivatized with the following steps. First, a 10 μL of methoxyamine hydrochloride in pyridine (40 mg/mL) was added to each sample and then shaken at 30°C for 90 min. Then 90 μL of N-methyl-N-(trimethylsilyl) trifluoroacetamide (MSTFA, Sigma-Aldrich) was added for trimethylsilylation. C8–C30 fatty acid methyl esters (FAMEs) were added as internal standard for retention time correction. Then samples were shaken for 30 min at 37°C. These derivatized samples were subjected to GC-TOFMS (Leco Pegasus IV) fitted with automatic liner exchange-cold injection (Gerstel). Agilent 6890 GC is equipped with a Gerstel automatic liner exchange system (ALEX) that includes a multipurpose sample (MPS2) dual rail, and a Gerstel CIS cold injection system (Gerstel, Muehlheim, Germany) with temperature program as follows: 50°C to 275°C final temperature at a rate of 12 °C/s and hold for 3 min. Injection volume is 0.5 μL with 10 μL/s injection speed on a splitless injector with purge time of 25 s. Liner (Gerstel #011711-010-00) is changed after every 10 samples using the Maestro1 Gerstel software vs. 1.1.4.18. Before and after each injection, the 10 μL injection syringe is washed three times with 10 μL ethyl acetate. A 30 m long, 0.25 mm i.d. Rtx-5Sil MS column (0.25 μm 95% dimethyl 5% diphenyl polysiloxane film) with additional 10 m integrated guard column was used (Restek, Bellefonte PA). 99.9999% pure helium with built-in purifier (Airgas, Radnor PA) was set at constant flow of 1 mL/min. The oven temperature was held constant at 50°C for 1 min and then ramped at 20°C/min to 330°C at which it was held constant for 5 min. A Leco Pegasus IV time of flight mass spectrometer was controlled by the Leco ChromaTOF software vs. 2.32 (St. Joseph, MI). The transfer line temperature between gas chromatograph and mass spectrometer was set to 280°C. Electron impact ionization at 70V is employed with an ion source temperature of 250°C. Acquisition rate was 17 spectra/s, with a scan mass range of 85-500 Da.

### Raw data files are preprocessed directly after data acquisition and stored as ChromaTOF-specific *.peg files, as generic *.txt result files and additionally as generic ANDI MS *.cdf files. ChromaTOF vs. 2.32 is used for data preprocessing without smoothing, 3 s peak width, baseline subtraction just above the noise level, and automatic mass spectral deconvolution and peak detection at signal/noise (s/n) levels of 5:1 throughout the chromatogram. Apex masses are reported for use in the BinBase algorithm. Result *.txt files are exported to a data server with absolute spectra intensities and further processed by a filtering algorithm implemented in the metabolomics BinBase database. The BinBase algorithm (rtx5) used the following settings: validity of chromatogram (10^7^ counts/s), unbiased retention index marker detection (MS similarity > 800, validity of intensity range for high m/z marker ions), retention index calculation by 5th order polynomial regression. Spectra are cut to 5% base peak abundance and matched to database entries from most to least abundant spectra using the following matching filters: retention index window ± 2,000 units (equivalent to about ± 2 s retention time), validation of unique ions and apex masses (unique ion must be included in apexing masses and present at > 3% of base peak abundance), mass spectrum similarity must fit criteria dependent on peak purity and s/n ratios and a final isomer filter. Failed spectra are automatically entered as new database entries if s/n >25, purity 80%. All thresholds reflect settings for ChromaTOF v. 4.0. Quantification is reported as peak height using the unique ion as default, unless a different quantification ion is manually set in the BinBase administration software BinView. A quantification report table is produced for all database entries that are positively detected in more than 10% of the samples of a study design class (as defined in the miniX database) for unidentified metabolites. A subsequent post-processing module is employed to automatically replace missing values from the *.cdf files. Replaced values are labeled as ‘low confidence’ by color coding, and for each metabolite, the number of high-confidence peak detections is recorded as well as the ratio of the average height of replaced values to high-confidence peak detections. These ratios and numbers are used for manual curation of automatic report data sets to data sets released for submission.

**Proteomics method :**

Each FFPE slide was incubated on 60° C for 45 min. Then, on room temperature, slides were washed with xylene (twice), ethanol (twice with100% and single time with 85% and 70%) and finally with deionized pure water. A 3 mm section were dissected from the slides and transferred into a vial with a buffer D (150 mM NaCl, 10 mM Tris-HCl pH 7.2, 2% SDS, 1% Triton X-100, 1% sodium deoxycholate, 5mM EDTA). After centrifugation at 10000 rcf for 1 min, samples were heated at 94°C for 30 min and 60 °C for 3 h.  Proteins were precipitated by adding chloroform/methanol in the samples and centrifuging them on 14000 rcf. Supernatant was discarded and protein pellet was dried in a speed-vac. The dried pellets were dissolved into 60 μL of 50 mM ammonium biocarbonate and were incubated for 10 min in a sonic bath. Samples were digested with 1 μg of trypsin for overnight at 37°C. Another 0.5 μg trypsin was added next morning and incubated for 3 more hours. Next, the samples were treated by a detergent removal kit (Pierce Biotechnology, IL, USA).

Digested peptides were analyzed by LC-MS/MS on a Thermo-Electron LTQ mass spectrometer, Michrom Paradigm LC and CTC Pal autosampler.  Peptides were directly loaded onto a Agilent ZORBAX 300SB C18, reversed phase trap cartridge, which, after loading, was switched in-line with a Michrom Magic C18 AQ 200 μm x 150 mm C18 column connected to the Thermo-Electron LTQ iontrap mass spectrometer through a Michrom Advance Plug and Play nano-spray source.  The nano-LC column (Michrom 3 µ 200 Å MAGIC C18AQ 200µm x 150 mm) was used with a 90 min gradient and a binary buffer system with buffer A composition 0.1% formic acid in water and buffer B composition 100% acetonitrile (gradient: 2-35% buffer B in 85 min, 35-80% buffer B in 23 min, 80% buffer B for 1 min, 80-2% buffer B in 1 min, 2% buffer B for 9 min) at a flow rate of 2 μL/min for the maximum separation of tryptic peptides.  The MS/MS spectra were acquired using a top 10 method, where the top 10 ions in the MS scan were subjected to automated low energy CID. An MS survey scan was obtained for the m/z range 375-1400, and MS/MS spectra were acquired using for the three most intense ions from the survey scan.  An isolation mass window of 2 Da was for the precursor ion selection, and a normalized collision energy of 35% was used for the fragmentation. A 2 min duration was used for the dynamic exclusion. These methods were used for the initial protein identifications, and not the posttranslational modification (PTM) analysis.

Tandem mass spectra were extracted with Xcalibur version 2.0.7. All MS/MS samples were analyzed using X! Tandem (The GPM, thegpm.org; version TORNADO (2010.01.01.4)). X! Tandem was set up to search the Uniprot Human Complete proteomics database supplemented with non-human common laboratory contaminants (<http://www.thegpm.org/cRAP/>)  (version August 4, 2010: 44512 entries, including 22256 reverse sequences) assuming the digestion enzyme trypsin. X! Tandem was searched with a fragment ion mass tolerance of 0.40 Da and a parent ion tolerance of 1.8 Da. Lodoacetamide derivative of cysteine was specified in X! Tandem as a fixed modification. Deamidation of asparagine and glutamine, oxidation of methionine and tryptophan, sulphone of methionine, tryptophan oxidation to formylkynurenin of tryptophan and acetylation of the n-terminus were specified in X! Tandem as variable modifications.

Scaffold (version Scaffold_3_00_04, Proteome Software Inc., Portland, OR) was used to validate MS/MS based peptide and protein identifications. Peptide identifications were accepted if they could be established at greater than 90.0% probability as specified by the Peptide Prophet algorithm (12403597).  Protein identifications were accepted if they could be established at greater than 90.0% probability and contained at least 2 identified peptides. Protein probabilities were assigned by the Protein Prophet algorithm (14632076). Proteins that contained similar peptides and could not be differentiated based on MS/MS analysis alone were grouped to satisfy the principles of parsimony.  The peptide and protein decoy false discovery rates (FDR) were calculated using the Scaffold software (18052118).
